# The Emergence of Non-Linear Evolutionary Trade-offs and the Maintenance of Genetic Polymorphisms

**DOI:** 10.1101/2024.05.29.595890

**Authors:** Samuel V. Hulse, Emily L. Bruns

## Abstract

Evolutionary models of quantitative traits often assume trade-offs between beneficial and detrimental traits, requiring modelers to specify a function linking costs to benefits. The choice of trade-off function is often consequential; functions that assume diminishing returns (accelerating costs) typically lead to single equilibrium genotypes, while decelerating costs often lead to evolutionary branching. Despite their importance, we still lack a strong theoretical foundation to base the choice of trade-off function. To address this gap, we explore how trade-off functions can emerge from the genetic architecture of a quantitative trait. We developed a multi-locus model of disease resistance, assuming each locus had random antagonistic pleiotropic effects on resistance and fecundity. We used this model to generate genotype landscapes and explored how additive versus epistatic genetic architectures influenced the shape of the trade-off function. Regardless of epistasis, our model consistently led to accelerating costs. We then used our genotype landscapes to build an evolutionary model of disease resistance. Unlike other models with accelerating costs, our approach often led to genetic polymorphisms at equilibrium. Our results suggest that accelerating costs are a strong null model for evolutionary trade-offs and that the eco-evolutionary conditions required for polymorphism may be more nuanced than previously believed.

## 1. Introduction

From life-history to foraging to disease resistance, genetic trade-offs are at the heart of many questions in evolutionary biology. In mathematical models, trade-offs between beneficial and deleterious traits are often necessary to maintain balancing selection [1]. Without an intrinsic downside, there is nothing preventing quantitative traits from evolving towards their maximum. For example, models of disease resistance typically assume the evolution of increased resistance carries a cost to either host mortality or fecundity [2,3]. Such trade-offs could emerge from either physiological constraints, or pleiotropic effects of the mutations affecting the focal trait.

Theoretical models of evolutionary processes have shown that particular assumptions about the shape of trade-off function, or how one quantitative trait scales with another can have major implications for evolutionary outcomes [3–7]. Disease resistance is particularly emblematic of trade-off function dependent evolution: Boots and Haraguchi [3] found when fecundity costs scale faster than resistance benefits (referred to as accelerating, or convex cost functions), evolution favours a single intermediate host genotype, whereas decelerating (also referred to as concave) costs lead to the coexistence of resistant and susceptible hosts. Accelerating costs leading to a single optimal genotype while decelerating costs lead to genetic polymorphisms is common outcome of models of quantitative traits with ecological feedbacks [5]. Similar patterns have been shown in predator behaviour models [8,9], life-history evolution models [10] and disease resistance evolution models [3,4]. The shape of trade-off functions has also been shown to determine evolutionary outcomes *in vivo*. By manipulating fecundity-survival trade-offs in *Escherichia coli*, Maharjan et al. were able to experimentally validate the results of theoretical models showing that changes in the shape of trade-off functions can indeed determine evolutionary outcomes [7].

Despite the abundance of evidence demonstrating the importance of trade-off functions, we understand their consequences far more than the biological processes that shape trade-off functions. While trade-off functions depict genotypic variation as a one-to-one relationship between quantitative traits, natural variation is two-dimensional. To address this, the concept of the Pareto front is a useful bridge [11]. The Pareto front is defined as the set of all phenotypes, such that improving performance in one trait can only be accomplished through a decrease in performance in another trait. For example, if we consider a trade-off between disease resistance and fecundity, the Pareto front represents the most fecund phenotypes for each level of resistance (Fig 1). Theory predicts that evolution should select for genotypes close to the Pareto front, with genetic polymorphism oriented along the front [12]. Mapping the curvature of the Pareto front can be used as a strategy to identify trade-off functions [13–16], thus understanding how genetic factors shape the Pareto front could be valuable for understanding genetic trade-offs.

**Figure 1:**
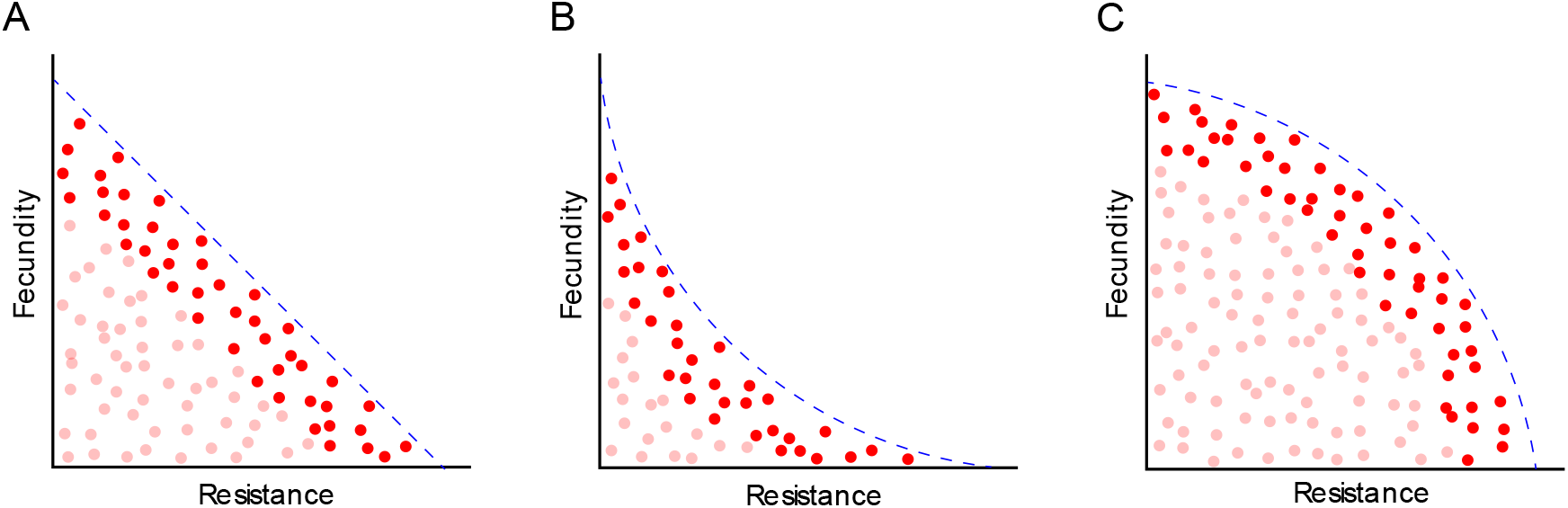
Schematic of how Pareto Fronts can define cost functions for (A): linear costs, (B): decelerating costs and (C): accelerating costs. Each dot represents a host genotype. The lighter red dots represent possible genotypes that would be removed by selection. The dashed blue line represents the Pareto front, where the phenotype space beyond is inaccessible to evolution.

If we assume a set of pleiotropic alleles have independent, additive contributions to a beneficial and detrimental trait, then low levels of the beneficial trait should be achievable using only the most cost-effective alleles. However, this might not be possible for higher levels of the beneficial trait, meaning evolution must have to rely on costlier alleles. This is one mechanism that could produce accelerating costs, although this prediction relies on strongly simplifying genetic assumptions, principally the absence of epistatic interactions between loci. If beneficial epistatic interactions between multiple alleles are only realized once multiple pleiotropic alleles are fixed, the benefits of subsequent mutations could be magnified, leading to decelerating costs.

To test the prediction that strictly additive genetics produces accelerating costs, while epistasis could produce decelerating costs, we developed an allelic model of the evolution of quantitative disease resistance. We assumed that disease resistance was determined by a series of discrete haploid loci, where each locus can have two possible alleles: one with antagonistic pleiotropic effects on fecundity and host resistance and one with no effects on either. With this framework, we generated genotype distributions and investigated the degree to which epistasis can influence the shape of the Pareto front. Next, we incorporated our genotype distribution model into an evolutionary model of disease resistance, where mutation allows hosts to move between genotypes. With this model, we asked whether epistatically induced changes in trade-off functions can result in a shift from a single dominant genotype to the maintenance of genetic polymorphism, mirroring patterns seen in previous models of quantitative disease resistance [3]. Unlike previous models [3,5], our approach requires no initial assumptions about trade-off functions.

## 2. Simulating Pareto Fronts

To simulate Pareto fronts, we developed a model of quantitative pathogen resistance (henceforth referred to as the discrete random loci model) which assumes that host resistance is determined by a fixed number, *n*, of haploid loci. Each locus has two possible alleles: a neutral allele which has no effect on the host phenotype, and an active allele which pleiotropically increases host resistance (benefits) and reduces host fecundity (costs). Given a sample of allelic effects, we can then define the set *G* of all possible genotypes as *G* = {0,1}^*n*^, with each genotype ***g***_***i***_ **∈ *G*** being a vector of length *n*. For each locus, a 0 represents the neutral allele while a 1 represents the active allele. This process can be thought of as flipping switches on a panel with *n* different switches, where each combination of switch positions produces a unique genotype. For each locus, we assumed that the active allele has costs and benefits sampled from a random exponential distribution. We define the resistance effect vector, ***r*** by *r*_*i*_∼ Exp(*λ*_*b*_) and the fecundity cost vector ***c*** by *c*_*i*_ ∼ Exp(*λ*_*c*_ ), where *i* ≤ *n*, λ_*c*_ represents the cost variance and λ_*b*_ represents the benefit variance. We initially assumed active alleles at multiple loci had additive effects for both resistance and fecundity. With this assumption, we define disease transmission, 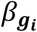 for a given genotype *g*_*i*_ as the normalized sum of all the active alleles for that genotype (Eqn. 1, subtracted from 1 to convert resistance to transmission). This value is then multiplied by *β*_0_, the baseline level of transmission.

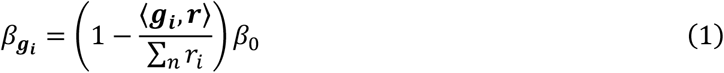

With this normalization, 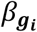 ranges from 0 to *β*_0_. The total fecundity cost of each genotype, 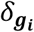 are defined similarly, where fecundity is normalized to range from 0.2 (the baseline deathrate, see Section 3 below) to 1.

Beyond purely additive allele interactions, we also explored how non-additive epistasis can influence the shape of the Pareto front. Here, a random subset of all active allele pairs is considered to have an epistatic interaction. For every interacting pair of alleles, epistasis either increases or decreases the combined effect of both alleles on the host resistance. We considered two forms of epistasis: first-order, where pairs of alleles have an epistatic interaction and second-order, where triplets of alleles have an epistatic interaction. There are 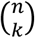 unique loci pairs, which could possibly have an epistatic interaction, where *n* is the number of loci and *k* = 2 for first-order epistasis or *k* = 3 for second-order epistasis. We randomly assigned a fixed proportion of these pairs and triples. We then assigned each pair an epistatic interaction with probability *p*_1_ for first-order epistasis and *p*_2_ for second-order epistasis. Next, we modified the cumulative effect of each loci pair (*i, j)* on resistance to *θ*_1_*r*_*i*_ + *r*_*j*_ where 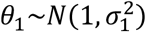. We implemented second-order epistasis similarly, by setting the cumulative effect of three given alleles to *θ*_2_*r*_*i*_ + *r*_*j*_ + *r*_*k*_ where 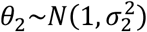. For all simulations with epistasis, the normalization step occurs after the epistatic effects are introduced.

We ran three series of simulations: one with no epistasis (Fig. 2A), one with only first-order epistasis (Fig. 2B), and one with both first and second order epistasis (Fig. 2C). For all simulations, we set the number of loci, *n*, to 9. Based on GWAS studies, this is a small but plausible number of loci [17–19]. Alternative models with either 5 or 13 loci did not have a qualitatively different effect on the Parent front (Fig. S1). For each epistasis treatment, we ran 100 random instantiations, and computed the Pareto front as the average optimal fecundity for every level of resistance (see the orange line, Fig 2A-C for an example of a single instantiation, Fig 2D-E for the average). Regardless of epistasis, the resulting Pareto had clear accelerating costs (Fig 2). However, with second-order epistasis, the trade-off function had a reduced curvature relative to other scenarios (Fig. 2C).

**Figure 2:**
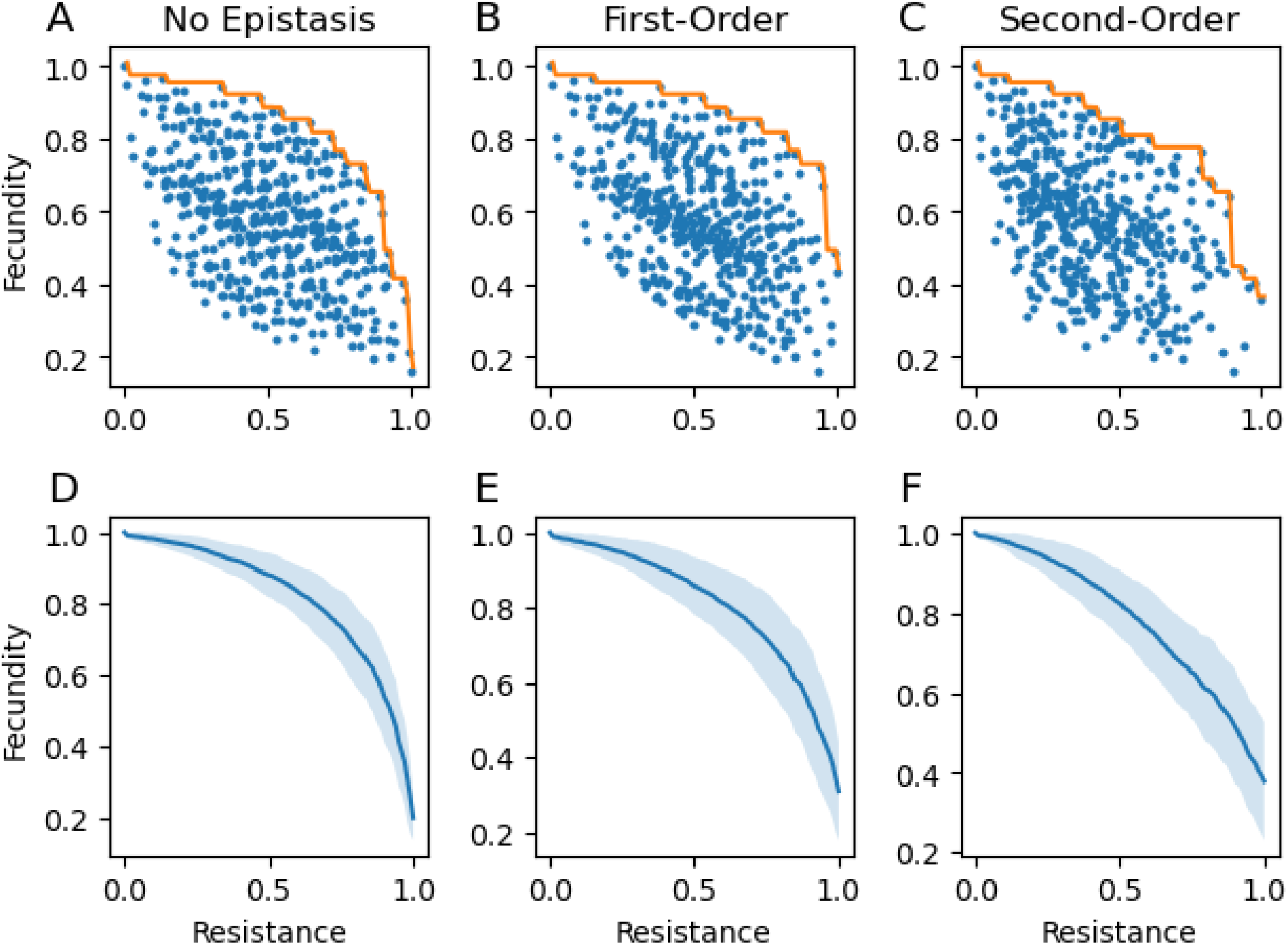
Distributions of genotype resistances and costs. A-C: Genotype distributions showing the fecundity and resistance level for all host genotypes for an instantiation with no epistasis (A), first-order epistasis (B), and first and second order epistasis (C). The clustering observed here is a byproduct of the additive loci: each additional locus effectively copies and shifts the distribution without it, leading to the observed patchiness when individual loci have large effects. For each, the orange line represents the Pareto front. D-F: Simulated Pareto front averaged over 100 instantiations for no epistasis (D), first-order epistasis (E), and first and second order epistasis (F). The light blue region represents values within one standard deviation of the average fecundity for a particular level of resistance. Parameters: λ_*b*_ = 0.1, λ_*c*_ = 0.1, *p*_1_ = 0.3, *p*_2_ = 0.3,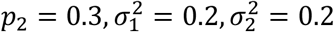.

## 3. Evolutionary Dynamics

To determine whether randomly generated genotype distributions drive similar evolutionary outcomes to standard models with trade-off functions, we built an evolutionary model on top of our randomly generated genotypes distributions. Given that we found accelerating costs when generating Pareto fronts, we predicted that this model would produce a single equilibrium genotype. Our model uses time-separated mutation and selection steps, similar to adaptive dynamics [20]. In classic adaptive dynamics models, mutations are introduced into populations at equilibrium, and this process is iterated until a final evolutionary equilibrium is reached. While adaptive dynamics models assume that new mutants differ from their parental generation by a small phenotypic value given by a trade-off function, our implementation assumes that new mutants differ from their progenitors by a single allele at a given locus. We used the discrete random loci model as the basis for the phenotype of each genotype, where a single mutation does not necessarily correspond to a small phenotypic change. Furthermore, instead of assuming a smooth trade-off function, trade-offs are generated by a random process and are inherently non-smooth.

For a given instantiation of random of allelic effects, each simulation begins with 100 uninfected hosts from the completely susceptible genotype, (*g*_0_ = (0, …, 0)) and 10 infected hosts. We assumed that hosts reproduce asexually. Furthermore, we assume a sterilizing, density-dependent disease without recovery, such that infection results in a total loss of fecundity without induced mortality. We then computed numerical solutions, from *t* = 0 to *t* = 1000, so that the hosts can reach the ecological equilibrium. At this point, we introduced mutation by taking 5% of all extant hosts and reassigning them to genotypes which differ from their progenitors by one allele. Analogous to adaptive dynamics, we then ran the simulation to ecological equilibrium again, and iteratively introduced new mutations. We ran the simulations for a total of 15 mutational iterations to reach evolutionary equilibrium, using the same genotype distribution parameters in Fig. 2. Since any two genotypes can differ by at most *n* loci, *n* mutational steps are sufficient for all possible genotypes to be reached. Since the shortest path to a particular genotype might not be evolutionary feasible, we include extra mutational iterations to allow for evolutionary equilibrium. The equations governing these dynamics are given below (Eqns. 2-3).

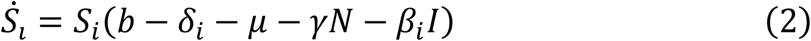

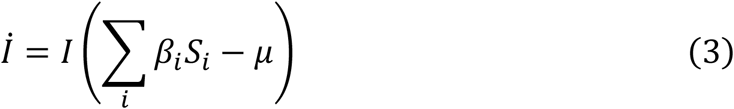

Here, *S*_*i*_ denotes the abundance of uninfected host genotype *i*, and *I* denotes the number of infected hosts and *N* represents the total number of hosts, both susceptible and infected. The host resistance and costs of resistance are given by *β*_*i*_ and *δ*_*i*_ respectively. Demographics are controlled by the birthrate, *b* the deathrate, *μ*, and the coefficient of density-dependent growth, *γ*. We considered three cases: no epistasis, first-order epistasis, and first and second-order epistasis (Fig 3).

**Figure 3:**
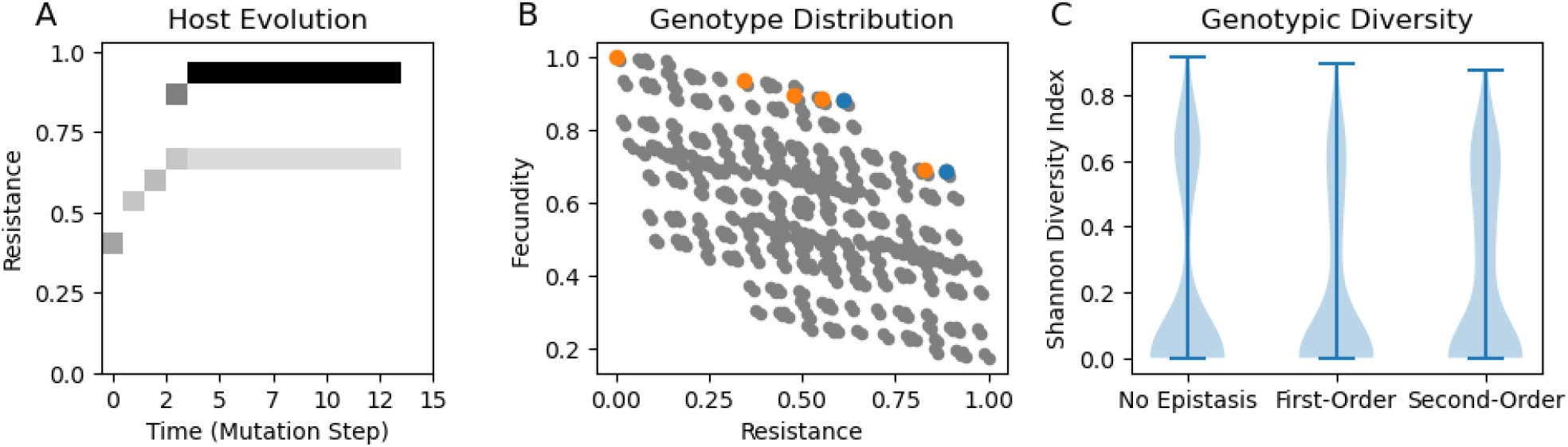
Evolutionary dynamics of disease resistance, using the discrete random loci model A: Phenotypic changes in resistance over the course of a simulation with no epistasis. B: Distribution of host genotypes with genotypes that had large populations at some point in evolutionary time in orange. Blue dots represent genotypes present at the end of the simulation, while orange dots are genotypes that were present at previous ecological equilibrium but not the final equilibrium. Panels A and B correspond to the same simulation. C: Equilibrium genetic diversity across simulations with 100 different allelic instantiations for each epistasis treatment. There was no significant difference in the Shannon diversity across treatments. Parameters: *β*_0_ = 0.005, *μ* = 0.2, *γ* = 0.001, the parameters for the trait distributions are the same as in Fig. 2.

To test whether epistasis affected equilibrium host genetic diversity, we ran 100 simulations for each epistasis treatment. We then calculated the host genetic diversity at equilibrium using the Shannon index, *H*, where *H* = − ∑_*i*_ *p*_*i*_ *ln*(*p*_*i*_ ), with *p*_*i*_ being the proportion of each genotype. Here, *H* = 0 indicates a monomorphic population, and *H* > 0 indicates a polymorphic population. As most simulations resulted in either one or two host genotypes, we used a non-parametric Kruskal-Wallis test to test whether the epistasis treatment produced significant differences in host diversity.

We expected diversity would be lowest in the purely additive model, since the Pareto front in the model was strongly accelerating, and in classic adaptive dynamics models only decelerating cost functions lead to stable polymorphisms. However, we found that certain model instantiations were able to maintain polymorphisms, even for purely additive models (Fig 3A-B). For simulations without epistasis, 30% had a polymorphism with at least 2 genotypes having equilibrium abundance greater than 5 (to ensure polymorphisms were not solely maintained by new mutants), while 37% were polymorphic for first-order epistasis and 43% for second-order epistasis. Epistasis did not significantly affect the host’s equilibrium genetic diversity (p = 0.54). We did not observe these polymorphisms when we ran adaptive dynamics simulations with equivalent parameters (Fig. S2).

## 4. Discussion

The discrete random loci model demonstrates how selection acting on a random assortment of mutations can generate non-linear cost functions. Our approach bridges fitness landscape models, such as the *N* − *k* model [21], and adaptive dynamics models [20] to test how genetic processes can define trade-off functions and enable polymorphism via ecological feedbacks. Our first key result is that accelerating cost function can emerge naturally from a process of random pleiotropic mutations followed by selection. We found that cost curves were most strongly accelerating when loci were purely additive. Epistasis could only blunt this trend but could not produce linear or decelerating costs. Our second key result is that allowing for a multilocus mutational process instead of a fixed trade-off function changes evolutionary predictions, resulting in more polymorphic outcomes. In single locus adaptive dynamics models, accelerating cost functions generally lead to stable, monomorphic populations [5]. However, we found that even though our mutation model generated Pareto fronts with an accelerating cost curve, the evolution and maintenance of stable polymorphism was common. This result contradicts the findings of classical adaptive dynamics models [3,5], suggesting that other eco-ecological factors beyond the shape of trade-off functions can drive genetic polymorphisms.

While some studies have demonstrated accelerating costs in experimental evolution studies [35–37], quantifying the relationship between traits and their costs is difficult. Costs can manifest in many different ways, potentially via specific ecological contexts [38], meaning that recreating the context in which costs manifest can be impractical if not impossible. For disease resistance, detecting any costs can be difficult, yet alone mapping costs to resistance levels with sufficient resolution to define a cost curve [39]. Despite this uncertainty, accelerating costs are a common assumption in adaptive dynamics models [5]. Our results suggest that this is a reasonable null model for trade-offs between quantitative traits, while decelerating costs might require more justification.

Contrary to our initial predictions, our model resulted in accelerating costs even with second-order epistasis. While decelerating costs might be more likely with third or even fourth order epistasis, such interactions are plausible but likely less frequent [22]. Decelerating costs might also emerge from evolvability constraints. In this case, even when the Pareto depicts accelerating costs, evolution may be unable able to track the front, instead following a path of decelerating costs. For example, the initial cost of evolutionary innovations may be reduced by compensatory mutations which can only emerge later [23]. Evolutionary trajectories may also depend on stepwise mutations at a single locus [24], as well as recombination, which are not included in our model. These simplifying assumptions in our model make it easier for evolution to reach all genotypes, potentially removing mechanisms that lead to a broader range of trade-off functions. Decelerating costs could also result from physiological constraints, such as allometric scaling laws. However, for a trait like quantitative resistance, it is not clear that such physiological constraints would have a greater role in defining trade-offs than additive genetic variance. Our evolutionary model of disease resistance differs from traditional adaptive dynamics approaches in several important ways. First, individual mutations do not necessarily result in small phenotypic changes. First, similar polymorphisms can emerge from models with only two alleles at a single locus [2]. Antonovics and Thrall found that when one host genotype is highly resistant, it allows for the coexistence of more susceptible genotypes by reducing the prevalence of infection. Since our model has the potential for single alleles with large effects, the same mechanism as in single locus models could produce polymorphisms. Secondly, as the Pareto fronts generated from my model are non-smooth, small perturbations from a purely accelerating cost function may result in small regions where costs grow at a decelerating rate relative to resistance. Such deviations can be seen in Fig. 2, where each curve has areas where it does not reflect the overall accelerating cost pattern. Since trade-offs in vivo are unlikely to be perfectly smooth [29], locally decelerating costs could be a plausible mechanism for maintaining genetic diversity in natural systems. This influence of both the smoothness of the trade-off curve could be tested by introducing perturbations into trade-off functions in adaptive dynamics models known to produce a single continuously stable strategy to see if that strategy remains stable with perturbation. If small regions of decelerating costs are responsible for polymorphism in our model, then this should be reflected through adaptive dynamics as well.

With the advent of modern genomics, many assumptions of our model are increasingly testable [25]. Our assumption that allelic effects are exponentially distributed is supported by population genetics theory and GWAS studies [26–28]. While quantifying the frequency and magnitude of epistatic interactions between many loci is difficult, combinatorial approaches to mapping out fitness landscapes can be illuminating [29–31]. Depending on the trait and model system, the frequency of epistasis is highly variable [25,32]. Furthermore, studies in yeasts and bacteria have found that higher-order epistasis is nearly as prevalent as pairwise epistasis, and that epistatic interactions occur between roughly 10% of mutation triplets [22,33,34]. While the prevalence of epistatic interactions is highly species and phenotype dependent, what we do know suggests that our implementation is a reasonable first approach.

In natural populations, traits like quantitative pathogen resistance often have a high degree of genetic variability, thus theoretical models must reflect how this variation is maintained. Our model demonstrates that without very strong epistasis, or a clear physiological mechanism, accelerating costs might be the most realistic cost function for most evolutionary trade-offs. However, unlike models with smooth trade-off functions, our model shows that even these accelerating cost functions can lead to polymorphic outcomes. Stochastic, jagged trade-off functions may therefore be an important driver of genetic variation.

## Ethics

This work did not require ethical approval from a human subject or animal welfare committee.

## Data accessibility

The python code used to generate all data and figures in this article is available on GitHub at https://github.com/svhulse/cost-model.

## Declaration of AI use

We have not used AI-assisted technologies in creating this article.

## Authors’ contributions

S.V.H.: conceptualization, formal analysis, software, visualization, writing - original draft. E.L.B.: funding acquisition, supervision, writing – review and editing.

## Conflict of interest declaration

We declare we have no competing interests.

## Funding

This work was supported by a grant from the NIH (grant number R01GM140457) to Michael Hood and Emily Bruns.

## Acknowledgements

We would like to thank Janis Antonovics for his encouragement, feedback, and many helpful discussions.

## Notes

https://github.com/svhulse/cost-model

### Competing Interest Statement

The authors have declared no competing interest.

