## Supplemental Figures for "The Emergence of Non-Linear Evolutionary Trade-offs and the Maintenance of Genetic Polymorphisms"

### Simulations with Alternate Numbers of Loci

To ensure that our results were robust to different numbers of loci underlying disease resistance, we ran addition simulations with fewer (5) or more (13) loci. In both cases, we observed similar outcomes as we did in our model presented in the main text with 9 loci (Fig. S1).

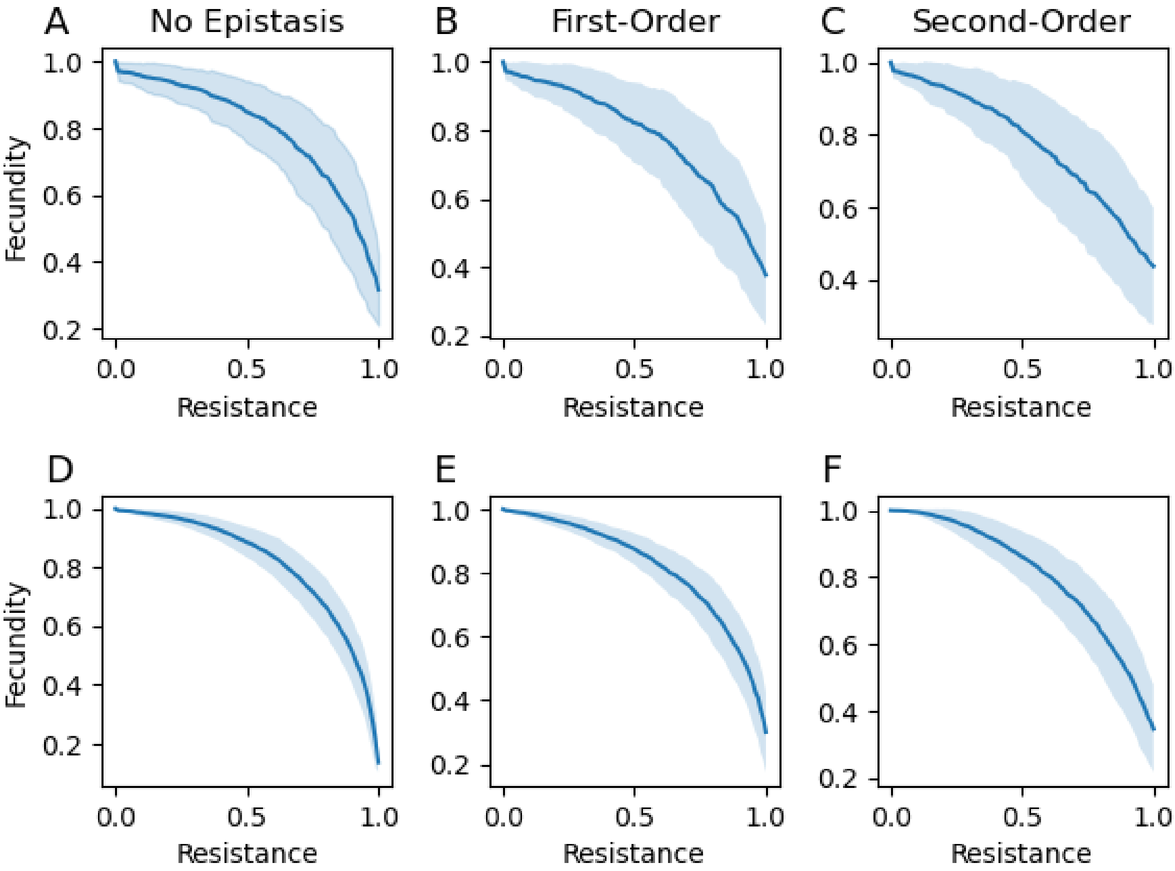


**Figure S1:** Average Pareto front for models with different numbers of loci. A-C: Five locus genotype distributions showing the fecundity and resistance level for all host genotypes for an instantiation with no epistasis (A), first-order epistasis (B), and first and second order epistasis (C). D-E: Thirteen locus genotype distributions showing the fecundity and resistance level for all host genotypes for an instantiation with no epistasis (D), first-order epistasis (E), and first and second order epistasis (F).

### Adaptive Dynamics Simulations

To test whether the polymorphisms we observed with the discrete random loci model were due to the shape of the emergent accelerating trade-off curve, or other properties of the model, we made an adaptive dynamics model with similar parameters to the discrete random loci map. For a random instantiation of genotypes, we first computed the Pareto front. We then fitted a sixth-degree polynomial to the Pareto front as a smooth approximation of the Pareto front. With this function, we built an adaptive dynamics model with the same system of equations as we used for the dynamical model in the main text (Eqns. 2-3). For any simulations that would produce polymorphisms with the discrete random loci model we found that the adaptive dynamics model led to a monomorphic equilibrium (Fig. S2).

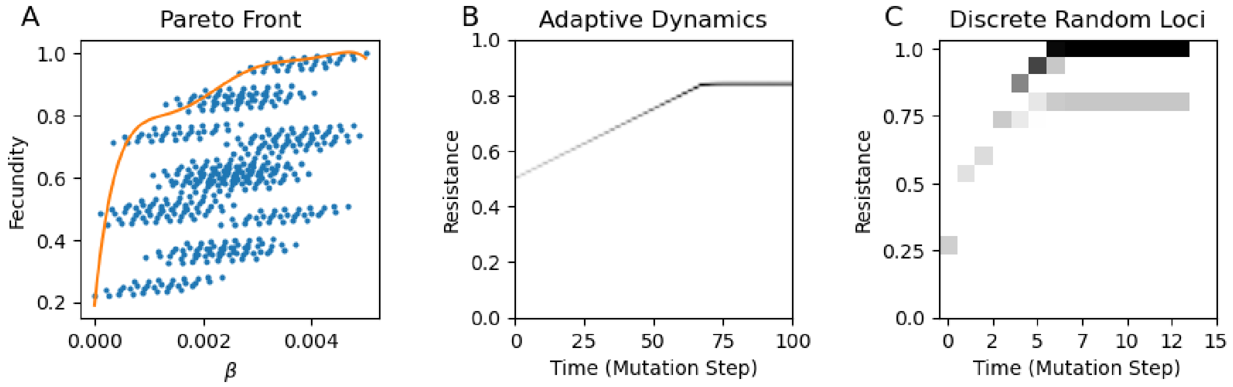


**Figure S2:** Adaptive dynamics model replicating the parameters of the discrete random loci model. (A): Randomly sampled genotype distribution with polynomial approximation of the pareto front. (B): Results from the adaptive dynamics model, where the cost function was taken from the polynomial approximation of the trade-off function, showing evolutionary change over time. (C): Results from the discrete random loci model, using the genotype distribution shown in (A).
